# Technical and biological sources of noise confound multiplexed enhancer AAV screening

**DOI:** 10.1101/2025.01.14.633018

**Authors:** Avery C. Hunker, John K. Mich, Naz Taskin, Amy Torkelson, Trangthanh Pham, Darren Bertagnolli, Anish Bhaswanth Chakka, Rushil Chakrabarty, Nicholas P. Donadio, Rebecca Ferrer, Molly Gasperini, Jeff Goldy, Junitta B. Guzman, Kelly Jin, Shannon Khem, Rana Kutsal, Jean-Benoît Lalanne, Refugio A. Martinez, Dakota Newman, Nick Pena, Nadiya V. Shapovalova, Natalie Weed, Thomas Zhou, Shenqin Yao, Jay Shendure, Kimberly A. Smith, Ed S. Lein, Bosiljka Tasic, Boaz P. Levi, Jonathan T. Ting

**Author notes:** Correspondence: Avery C. Hunker.

## Abstract

*Cis*-acting regulatory enhancer elements are valuable tools for gaining cell type-specific genetic access. Leveraging large chromatin accessibility atlases, putative enhancer sequences can be identified and deployed in adeno-associated virus (AAV) delivery platforms. However, a significant bottleneck in enhancer AAV discovery is charting their detailed expression patterns *in vivo*, a process that currently requires gold-standard one-by-one testing. Here we present a barcoded multiplex strategy for screening enhancer AAVs at cell type resolution using single cell RNA sequencing and taxonomy mapping. We executed a proof-of-concept study using a small pool of validated enhancer AAVs expressing in a variety of neuronal and non-neuronal cell types across the mouse brain. Unexpectedly, we encountered substantial technical and biological noise including chimeric packaging products, necessitating development of novel techniques to accurately deconvolve enhancer expression patterns. These results underscore the need for improved methods to mitigate noise and highlight the complexity of enhancer AAV biology *in vivo*.

## Main

Enhancers are *cis*-acting DNA elements that interact with transcription factors to increase the probability of endogenous gene expression. Depending on the chromatin state and transcription factors present, enhancer-driven gene expression *in vivo* can be highly cell type- and region-dependent. Recent advancements in single cell sequencing technologies have facilitated large-scale enhancer identification across cell classes, subclasses, and types through the integration of cell type classification with epigenomic discovery^1,2^. Machine learning algorithms trained on large genomic datasets can accurately predict enhancer activity based on sequence motifs, chromatin accessibility, and epigenetic modifications^3,4^. Importantly, these putative enhancer sequences are portable into synthetic contexts and can be packaged alongside a minimal promoter into enhancer AAVs which can drive cell type-specific gene expression^5^. These efforts have resulted in a large toolbox containing thousands of enhancer AAVs for targeting diverse cell types across the brain^6–12^.

To validate enhancer AAVs as functional tools in the brain, the most prominent and reliable method is laborious one-by-one testing. This involves pairing novel enhancer sequences with fluorescent reporters, individual AAV packaging and *in vivo* injections, and examining each enhancer independently for on-target cell type specificity and expression patterns. This screening approach requires an enormous amount of time, effort, and animals, making it challenging to perform duplicates or troubleshoot when enhancer AAVs exhibit unpredicted expression patterns. Parallelizing or multiplexing enhancer AAVs for validation would not only accelerate the screening process but may also provide avenues for retesting or optimizing enhancers with unexpected expression.

Massively parallel reporter assays (MPRAs) have been developed to enable simultaneous evaluation of thousands of putative enhancer sequences by detecting enhancer-driven mRNA transcripts^8,13–16^. However, most MPRAs screen hundreds of enhancers but lack the detection granularity necessary for calculating precise enhancer AAV specificity across cell types and extending these studies to single cell resolution has been difficult.

We devised a simple barcoded vector multiplex screen for enhancer AAV validation. We paired previously validated cell type-specific enhancers with small barcodes in the 3’ untranslated region (UTR), packaged the vectors into AAVs, injected them as a pool into mice, and used droplet-based RNA sequencing to identify enhancer-driven expression. Surprisingly, we observed substandard enhancer specificities which led us to investigate and identify numerous confounding variables that obscured true enhancer expression. Here we demonstrate weak AAV transcript detection combined with packaging chimeras (see also Lalanne et al., co-submitted) as well as enhancer AAV transcriptional crosstalk together cause high levels of technical and biological noise in a multiplex enhancer screen. We implemented novel techniques that helped to minimize background noise and improve barcode transcript detection but ultimately found we could not fully recapitulate the performance of individual enhancer testing. These findings highlight the inherent challenges of multiplex enhancer screening, emphasizing the importance of continued optimization to achieve more accurate and reliable enhancer AAV characterization.

## Results

### Design of a multiplex enhancer AAV validation pipeline in the mouse brain

Enhancer AAV validation is a multi-step process, involving cloning enhancer elements into reporter vectors, packaging into AAVs, injections into mice, and expression analysis (**Fig. 1a**). To establish a scalable workflow for robust enhancer AAV transcript recovery and cell type identification, we designed a strategy for multiplexed testing of enhancer specificity using barcoded vectors that allows each mouse to be injected with multiple enhancer AAVs (**Fig. 1b**).

**Figure 1:**
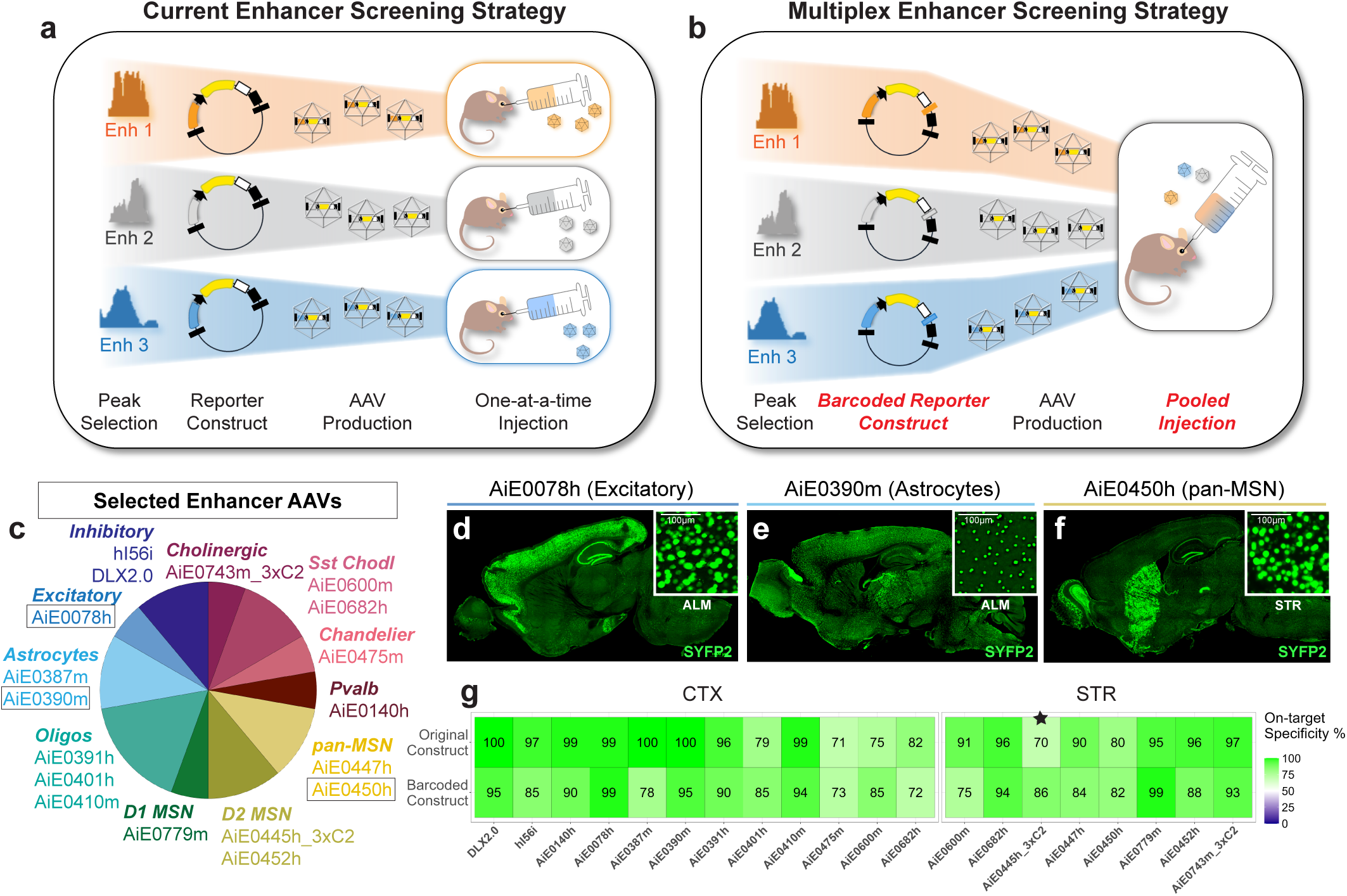
Multiplex screen design for rapid enhancer AAV characterization. **a)** Current one-by-one enhancer screening method. Enhancer AAVs are cloned, packaged, injected, and analyzed individually in mice for cell type specificity. **b)** Design of multiplex strategy. Barcoded reporter constructs permit pooling of enhancer AAVs for injection and simultaneous analysis of cell type specificity. **c)** Set of published enhancers used for validating multiplex screening method. Enhancers are grouped by their cell type specificities. Enhancers in boxes are highlighted in panels d-f. Size of pie slices corresponds to proportion of enhancers representing each cell type. **d-f)** Examples of nuclear SYFP2 fluorescence from barcoded enhancer AAVs injected RO individually into mice (d: pan-excitatory enhancer AiE0078h, e: astrocyte enhancer AiE0390m, f: pan-MSN enhancer AiE0450h). **g)** Comparison of enhancer AAV cell type specificities from the originally published construct (Mich et al 2021, Mich et al 2023, Ben-Simon et al 2024, Hunker, Wirthlin et al 2024) with barcoded constructs (n=1 per enhancer) used in this study in the cortex (CTX, left) or striatum (STR, right). Enhancer specificities for Sst Chodl enhancers AiE0600m and AiE0682h are reported and calculated for both brain regions. Reported specificity for AiE0445h_3xC2 is for the full-length version AiE0445h (indicated with a star).

To test our multiplex strategy, we selected eighteen previously published enhancers with high specificity for their intended cell type targets (greater than 70% on-target specificity)^2,6,7,10^ (**Fig. 1c-f**). This pool included five glial enhancers, with two astrocyte (AiE0387m and AiE0390m) and three oligodendrocyte (AiE0391h, AiE0401h, and AiE0410m) enhancers, and thirteen neuronal enhancers. Seven of the neuronal enhancers had regional expression patterns, with six labeling subpopulations of the striatum (pan-MSN enhancers AiE0450h, AiE0447h, and AiE0445h_3xC2; D2 MSN enhancer AiE0452h; D1 MSN enhancer AiE0779m; striatal cholinergic enhancer AiE0743m_3xC2) and one labeling chandelier cells, a subpopulation of parvalbumin interneurons in the cortex (AiE0475m). Three enhancers express broadly in neuronal classes, including two forebrain GABAergic enhancers (hDLXI56i and the optimized version, DLX2.0), and one pan-excitatory enhancer (AiE0078h). The last three enhancers labeled interneurons in multiple brain regions including inhibitory parvalbumin interneurons (AiE0140h) and Sst Chodl interneurons (AiE0682h and AiE0600m).

Each enhancer was paired with a unique 8 base pair (bp) barcode, subcloned into a nuclear reporter vector (SYFP2-H2B), and packaged individually into PHP.eB, an AAV capsid variant known to cross the blood-brain barrier (BBB) in C57BL6J mice^17^. We used immunohistochemistry to confirm enhancers driving the new barcoded nuclear construct closely matched the specificities of the originally published constructs containing cytosolic SYFP2 (**Fig. 1g** and **Fig. S1**). While all enhancers maintained >70% specificity for their intended cell type target, on-target specificities were on average slightly lower for the barcoded constructs (88 ± 8% compared to 92 ± 10%, mean ± standard deviation). The greater sensitivity of the barcoded nuclear reporter likely revealed low-level off-target expression that was not detected with the less-sensitive cytosolic SYFP2 reporter in earlier studies.

### Nuclear brain tissue preps are insufficient at capturing AAV transcripts in 10X Genomics RNA sequencing

We next sought to establish a cell isolation method that would permit unbiased cell type recovery (**Fig. 2a**). Both whole cell and nuclear isolations are used widely in RNA sequencing but offer different advantages^18^. Nuclear isolations are preferred to whole cell isolations because of their simple homogenization procedures and ability to utilize frozen tissues.

**Figure 2:**
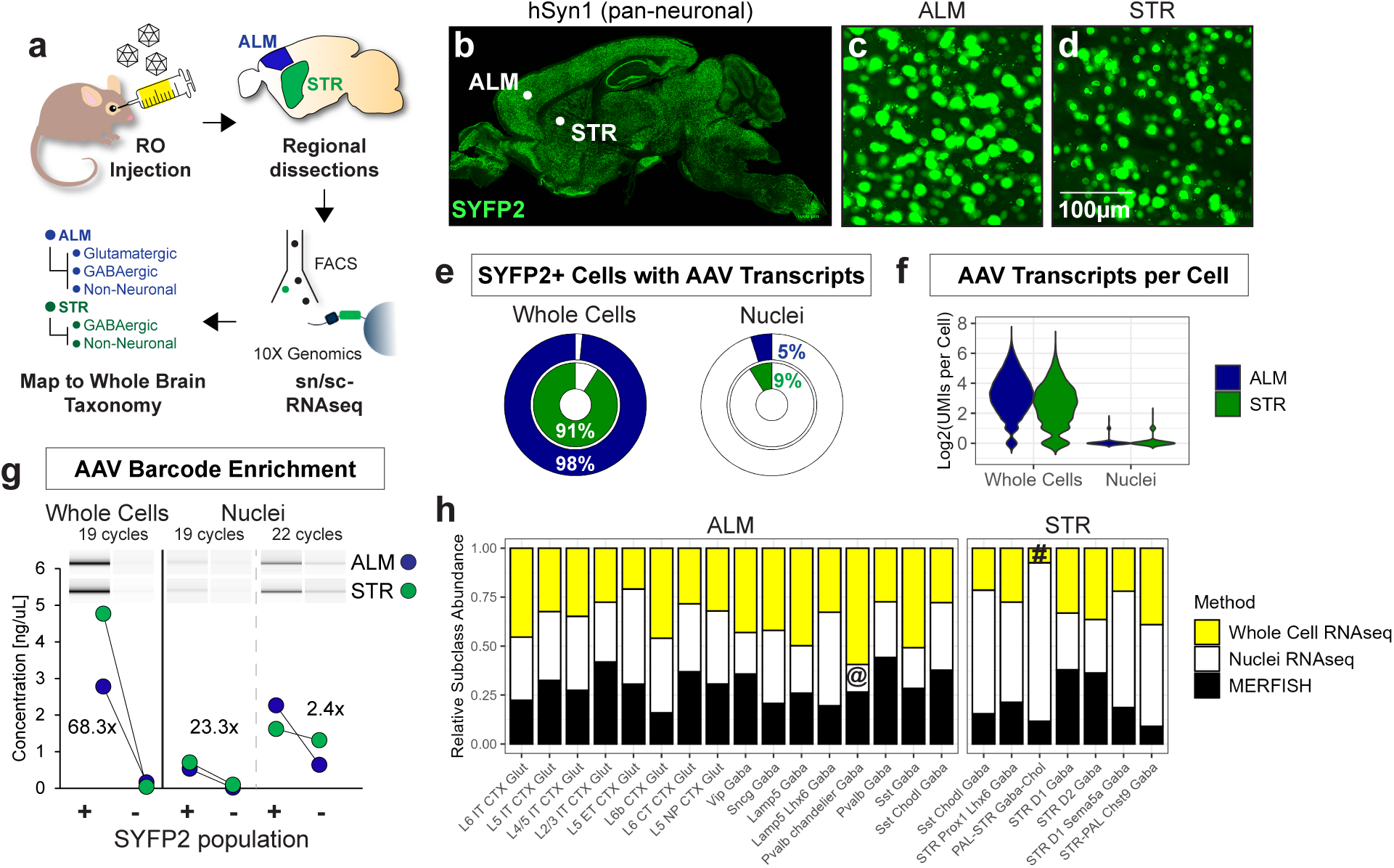
Robust AAV transcript detection is dependent on tissue prep method. **a)** Workflow for AAV screening by single whole cell or nucleus RNA sequencing. Cell type AAV expression is determined by mapping RNA sequencing reads to a mouse whole brain taxonomy. **b-d)** Images of brain wide nuclear SYFP2 fluorescence driven by pan-neuronal promoter hSyn1 with zoomed in views of ALM (c) and STR (d). **e)** AAV transcripts in ALM (blue outer ring) and STR (green inner ring) are detected in >90% of SYFP2-positive cells but <10% of SYFP2-positive nuclei. **f)** Number of unique AAV transcripts detected per cell or nucleus (UMIs) for ALM and STR. **g)** Comparison of enrichment strategy using custom AAV specific primers for barcoded transcripts in whole cells vs nuclei. Amplification from whole cell preps (left) result in reliable cDNA yields with minimal background, while same PCR cycling from nuclei results in low cDNA yields (middle). Increasing cycling in nuclei exposed ambient transcripts in the SYFP2-negative populations (right). Bioanalyzer virtual gel is of three separate gels with bands for ALM (top) and STR (bottom) for SYFP-positive and SYFP-negative cell populations. **h)** Comparison of cell type recovery in tissue prep methods to the cell type distribution in the published MERFISH mouse spatial dataset Zhuang ABCA-2 (https://portal.brain-map.org). MERFISH cell counts are from caudate putamen (CP) for STR and primary motor cortex (MOp) for ALM. # indicate subclasses underrepresented in cells; @ indicate subclasses underrepresented in nuclei.

We cloned the pan-neuronal promoter hSyn1 into the barcoded SYFP2-H2B screening vector and confirmed uniform SYFP2 expression throughout the brain (**Fig. 2b-d**). We generated suspensions of whole cells and nuclei from both the dorsal striatum (STR) and antero-lateral motor cortex (ALM), two brain regions with long projection neurons^18–20^. Both SYFP2-positive and SYFP2-negative cells and nuclei were FACS sorted and prepped for single cell/single nucleus RNA-seq (sc/snRNA-seq) using the 10X Genomics 3’ gene expression v3.1 platform. Cell types were identified by mapping sequenced transcriptomes to a published whole mouse brain taxonomy^21^.

We compared data from whole cells and nuclei and found significant differences in AAV transcript detection between methods (**Fig. 2e-f**). While approximately 10 AAV transcripts on average (determined by unique molecular identifiers, UMIs) were detected in >90% of SYFP2-positive whole cells, less than 10% of SYFP2-positive nuclei had any detectable transcripts (average of 1 AAV transcript/UMI per cell). We attempted to recover more AAV barcoded transcripts using an AAV-specific PCR from the gene expression (GEX) library cDNA (**Fig. 2g**). While 19 cycles provided a large and specific increase in AAV transcripts in the SYFP2-positive whole cell cDNA, the same PCR cycling from nuclear-prepared cDNA resulted in AAV transcript detection that was barely above background. Nuclear-prepped cDNA yielded AAV transcripts in SYFP2-negative samples at a level similar to SYFP2-positive samples when PCR cycles were increased to 22 cycles. These results suggest these AAV transcripts are depleted in nuclei, regardless of detection technique, and not due to technical limits.

We next compared ratios of recovered nuclei and whole cell types to the publicly available Allen Institute MERFISH mouse gene spatial dataset ^22^ (**Fig. 2h**). In general, all cell types were present across methods although we did note some small, insignificant variations in cell type proportions. Cortical chandelier cells (i.e. Pvalb Chandelier Gaba) and striatal cholinergic cells (i.e. PAL-STR Gaba-Chol) were slightly depleted in the nuclear and whole cell preps, respectively. While tissue preparation method substantially affects AAV transcript detection, we found no systematic effects on cell type recovery using cell or nucleus isolation methods. Thus, for further experimentation below we exclusively used whole cell preparations.

### Individual AAV packaging is required for multiplexed screening in the brain

With methods to detect barcodes developed, we designed a small multiplex using six AAVs for driving nuclear SYFP2 in distinct cell populations with little off-target expression^2,6,7^. We included AAVs containing four different enhancers (oligodendrocyte enhancer AiE0410m, astrocyte enhancer AiE0390m, forebrain GABAergic enhancer DLX2.0, and D1 MSN enhancer with striatum-only specificity AiE0779m) and two control promoters (hSyn1 and CMV) (**Fig. 3a**). All four enhancer AAVs included had >95% on-target specificity when injected individually (**Fig. 1g**).

**Figure 3:**
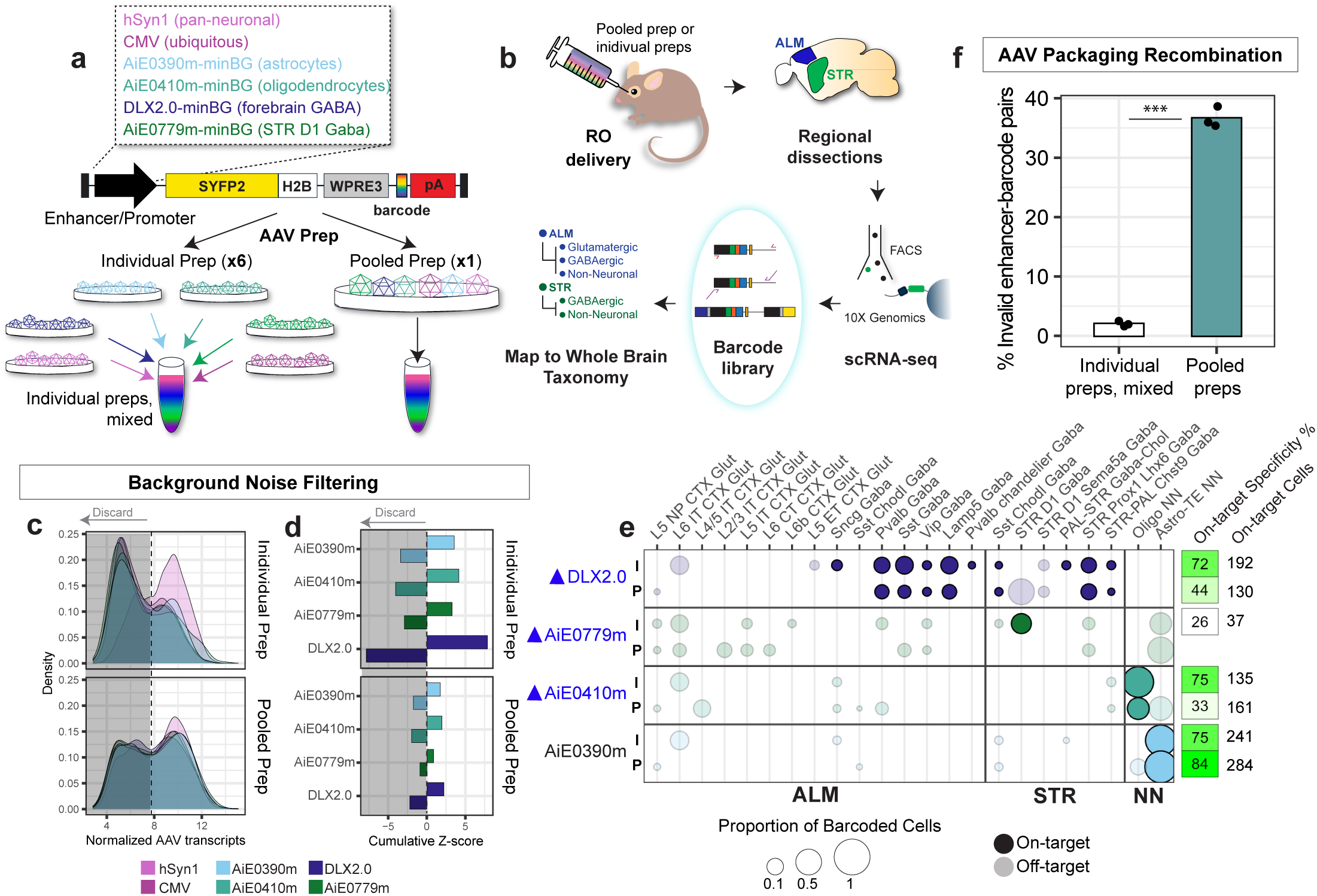
Diminished enhancer specificity and evidence of barcode swapping using pooled AAV packaging. **a)** Four enhancers (AiE0390m, AiE0410m, DLX2.0, and AiE0779m) along with two control promoters (CMV and hSyn1) were subcloned into barcoded constructs and packaged either separately as individual preps or combined at the plasmid level and packaged in a single pooled AAV prep. **b)** Workflow of experiment. Pooled AAV prep or six individual AAV preps mixed at equal proportions were injected RO into adult mice. SYFP2-positive cells from ALM and STR were sorted and prepped for scRNA-seq. Cell type AAV expression was determined by AAV barcode enrichment (barcode library) and mapping gene expression library to a mouse whole brain taxonomy. **c)** Distribution of transcripts per cell (UMIs) for each enhancer in individual (top) vs pooled (bottom) preps. Cells were discarded if normalized transcript UMIs were below the cutoff (dotted line). **d)** Cumulative Z-scores across enriched (positive) or depleted (negative) enhancer labeled subclasses in individual (top) vs pooled (bottom) preps. Enhancers from individual preps have higher enrichment Z-scores compared to pooled preps. **e)** Deconvolved enhancer specificities from individual (I, top) and pooled (P, bottom) preps. Dot size represents proportion of labeled cells for each enhancer. On-target cells are highlighted in full color and off-target cells are opaque. Dot colors represent enhancer cell type targets as denoted in A. On-target enhancer specificity percentages and number of on-target cells recovered are on the far right. Blue triangles indicate enhancers that lost specificity in pooled AAV prep. **f)** Long read sequencing of pooled and mixed individual AAV preps (n=3). Welch Two Sample t-test was performed to compare individual preps and pooled preps (t = −32.342, df = 2.1773, ***p-value = 0.0005813. The 95% confidence interval for the difference in means is [−0.3888412, −0.3035556]).

We tested whether we could increase screening efficiency by pooled packaging in which all six barcoded plasmids were mixed at equal ratios and packaged into a single AAV prep, similar to previous MPRAs^14^. Presence of all six barcodes was confirmed by deep sequencing the pooled plasmids, mixed individual AAV preps, or the single pooled AAV prep (**Fig. S2a**). We injected both pooled and mixed individual preps RO into mice separately and performed 10X scRNA-seq on whole cells isolated from the ALM and STR (**Fig. 3b** and **Fig. S2b**).

In contrast to hSyn1-driven transcript detection alone from Figure 2, enhancer AAV barcodes were much more difficult to capture. Despite sorting for SYFP2-positive cells, we only found enhancer AAV barcodes in approximately one third of cells in each of the gene expression (GEX) multiplex libraries (**Fig. S2c**). To boost AAV transcript detection, we generated a separate enhancer barcode library using PCR from the cDNA and found we were able not only to increase AAV transcript detection two-fold across cells, but also increase the number of enhancer barcodes per cell (**Fig. S2d**). Using presence of barcode alone as indication of enhancer expression, initial analysis suggested on-target enhancer specificities across both the GEX and enhancer barcode libraries were much lower than expected (<40% on-target cells compared to >95% by individual injection) for both the pooled and individual AAV preps (**Fig. S2e**).

We suspected the low specificity was caused by technical artifacts such as PCR chimeras, sequencing artifacts, or ambient RNA, which was supported by the enhancer barcode UMI bimodal distribution^31^ (**Fig. 3c**). The noise reduction algorithm Cellbender^23^ was unsuccessful in recovering expected enhancer specificities, suggesting ambient RNA and/or sequencing artifacts were not primary drivers of noise. Instead, we removed cells with enhancer barcode counts below the antimode of the bimodal distribution and then calculated Z-scores as a measure of enrichment for each cell type and discarded de-enriched cell populations (cell types with negative Z-scores) for each enhancer (**Fig. 3d**). Of note, we observed enhancers from the individual AAV preps had larger Z-scores and thus greater cell type enrichment than the same enhancers in the pooled prep, suggesting AAV preparation method may impact enhancer expression.

While all four enhancers labeled on-target cells, we found a large discrepancy in on-target specificity not only between the individual screening values and multiplex but also between AAV packaging methods (**Fig. 3e**). Three out of the four enhancer AAVs exhibited higher on-target cell type specificity in the mixed individual preps compared to the pooled preps (DLX2.0: 72% individual vs 44% pooled preps, AiE0779m: 26% individual vs 0% pooled preps, AiE0410m: 75% individual vs 33% pooled preps.). The astrocyte enhancer AiE0390m was the only enhancer with comparable specificities across both AAV prep methods (75% on-target in individual preps vs 84% on-target in pooled prep), possibly due to transduction biases of the AAV capsid PHP.eB. All enhancers across multiplexes had lower on-target specificities than when injected alone.

We speculated the low cell type enrichment and low on-target cell type specificities in the pooled AAV preps could be caused by recombination during packaging as all vectors contain largely similar open reading frames. We performed long read sequencing directly from AAV preps^24^ (service from Plasmidsaurus) and were surprised to find approximately 37% of reads contained mismatched enhancer-barcode pairings compared to only 2% in the mixed individual preps (**Fig. 3f**). Similar chimerism was observed across all three replicate packagings of the pooled six-plex library, and this result strongly agrees with independent experimentation (Lalanne et al., co-submitted). These findings reveal extensive barcode swapping during AAV pooled packaging that significantly compromises cell type specificity measurements from barcoded enhancer AAVs.

### AAV concatenation contributes to biological noise

Following transduction, AAVs can persist in the nucleus as concatemers^25–27^. While concatenation promotes long-term transgene expression, this configuration also can induce transcriptional crosstalk between enhancer AAVs, which occurs when an active enhancer from one vector drives expression of another linked vector ^28^. We hypothesized this biological phenomenon could contribute to aberrant enhancer barcode detection in the multiplexed experiment.

To test this, we obtained severe combined immunodeficiency (SCID; *Prkdc*^scid/scid^) mice which have loss of function of the DNA double-strand break repair enzyme Prkdc and have reduced AAV concatenation^25^. We injected all eighteen previously validated and individually packaged enhancer AAVs (**Fig. 1c**) plus two control promoters as a batch into either C57Bl6J or SCID mice. Deep sequencing of the 3’ UTR of the pooled viruses indicated similar representation of each barcode (**Fig. 4a**). SYFP2-positive whole cells were collected via FACS from the ALM and STR and processed for 10x sc-RNAseq using our established protocol (**Fig. 4b**).

**Figure 4:**
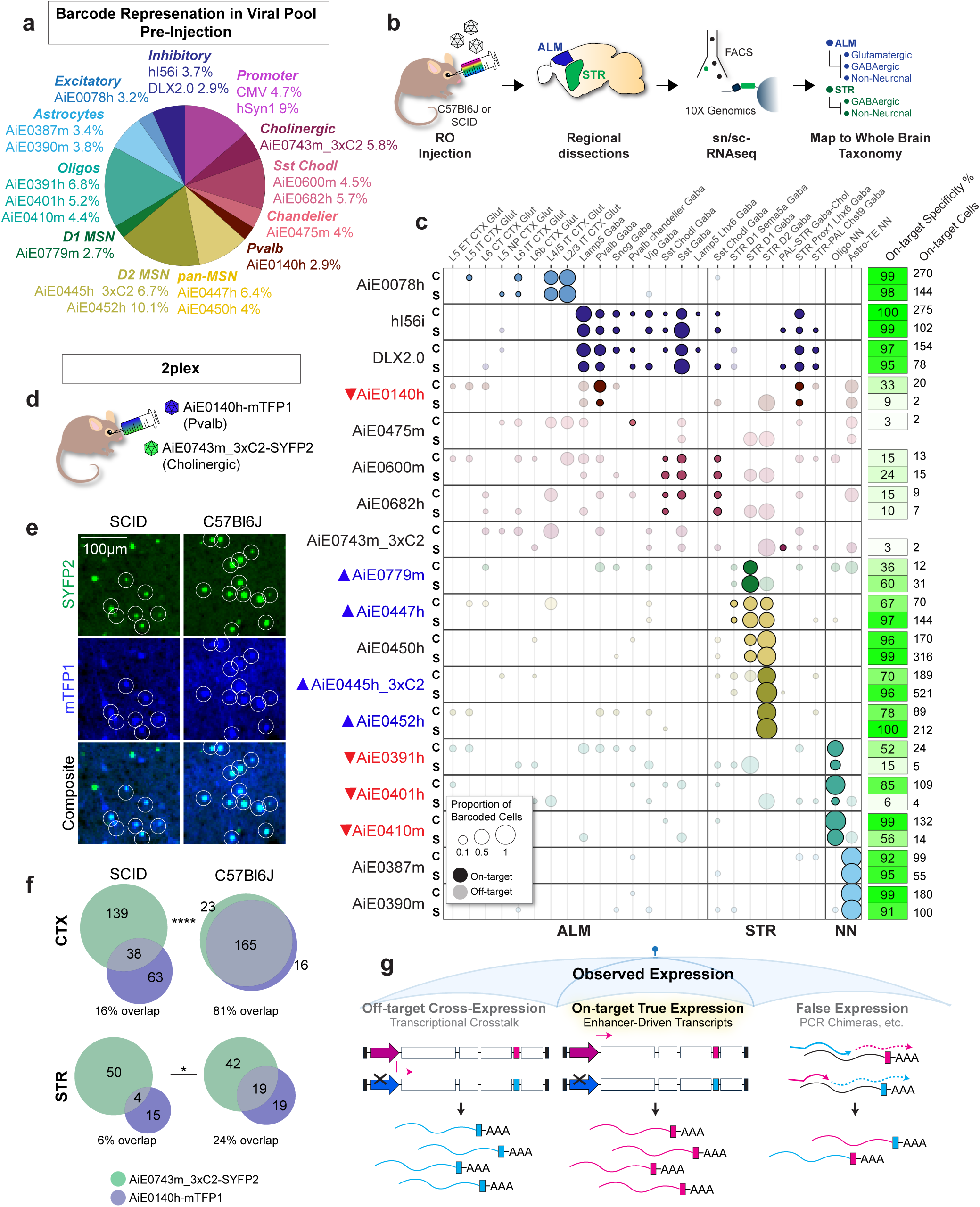
Multiplex screen in SCID mice attenuates biological noise in striatal cell types. **a)** Enhancers/promoters included in screen. Percentages are barcode representations in AAV pool pre-injection determined by Amplicon-EZ short read deep sequencing. **b)** Multiplex experiment workflow. C57Bl6J or SCID adult mice are injected RO with 20plex enhancer AAV pool. SYFP2-positive cells from ALM and STR are sorted, sequenced, and mapped to whole mouse brain taxonomy to determine cell identities. **c)** Deconvolved enhancer specificities from SCID (S, bottom) and C57Bl6J (C, top) mouse 20plex. Only labeled subclasses with positive Z-scores are shown and dot size represents proportion of labeled cells for each enhancer. Dot colors represent enhancer cell type targets as denoted in a. On-target enhancer specificity percentages and number of on-target cells recovered per enhancer are on the far right. Specificities for four striatal enhancers improved in SCID mice (blue triangles) while specificities for oligo enhancers and Pvalb enhancer AiE0140h decreased (red triangles). On-target cells are highlighted in full color and off-target cells are opaque. **d-f)** Two-plex pool containing striatal cholinergic enhancer AiE0743m_3xC2 driving SYFP2 and cortical Pvalb enhancer AiE0140h driving mTFP1 in both SCID and C57Bl6J mice (n=1 mouse per genotype). **d)** Pooled RO injection of enhancer AAVs. **e)** Native fluorescence from cortex with mTFP1+SYFP2+ cells circled. **f)** Venn diagram with quantification of mTFP1+ and SYFP2+ cells in the cortex (CTX) and striatum (STR) with number of cells counted as labels. Fisher’s Exact Test (two-sided) was used to evaluate enhancer overlap in SCID vs C57Bl6J (n=1 ROI per region). CTX: Odds ratio= 0.196 (95% CI: 0.128–0.296), with a p-value < 2.2e-16. STR: Odds ratio= 0.246 (95% CI: 0.058–0.789), with a p-value = 0.0116. **g)** Observed expression in multiplex screen is a combination of off-target cross-expression, on-target true expression, and false expression.

Similar to previous reports^25,28^, we observed a reduction in AAV-derived transcripts in the SCID mice, with the vast majority of enhancers (15/18, 83%) exhibiting less transcripts per cell compared to the C57Bl6J multiplex (**Fig. S3**). We also observed most enhancers (11/18 enhancers, 61%) illustrated similar specificities in both the C57Bl6J and SCID multiplex (**Fig. 4c**). The six enhancers that maintained high specificities (>90%) in both strains of mice included three enhancers with broad cell class specificities (pan-excitatory enhancer AiE0078h and both pan-inhibitory enhancers hI56i and DLX2.0), as well as one strong pan-MSN enhancer (AiE0450h) and both astrocyte enhancers (AiE0387m and AiE0390m). Interneuron enhancers were the least successful, with all five (Pvalb enhancer AiE0140h, Pvalb chandelier enhancer AiE0475m, Sst Chodl enhancers AiE0600m and AiE0682h, and striatal cholinergic enhancer AiE0743m_3xC2) labeling more off-target than on-target cells across both C57Bl6J and SCID multiplexes. This is likely because these are relatively rare cell types, each encompassing less than 1% of the reference taxonomy.

The remaining seven enhancers had substantially altered specificities between multiplex experiments. All MSN enhancer specificities were similar to individual screening methods (**Fig. 1g**) in the SCID mice, with both pan-MSN enhancers (AiE0447h and AiE0450h) and D2 MSN enhancer (AiE0445h_3xC2 and AiE0452h) reaching >95% specificity. The D1 MSN enhancer AiE0779m, although still below the individual testing, increased specificity to 60% for D1 MSNs in the SCID mice compared to 36% in C57Bl6J mice. In contrast, the on-target specificities for all three oligodendrocyte enhancers (AiE0391h, AiE0401h and AiE0410m) were much lower in the SCID mice. For example, despite having 85% specificity for oligodendrocytes by both individual injection and C57Bl6J multiplex, AiE0401h was only 6% specific in SCID mice. Additionally, the total number of barcoded oligodendrocytes collected was over 11-fold lower in the SCID multiplex (265 oligodendrocytes in C57Bl6J multiplex vs 23 oligodendrocytes in SCID multiplex).

Other than the oligodendrocyte enhancers, the cortical Pvalb interneuron enhancer AiE0140h also had reduced specificity and labeling of Pvalb interneurons in the SCID multiplex (9% on target vs 33% in C57Bl6J). To further investigate this, we co-injected AiE0140h driving the fluorescent protein mTFP1 and the striatal cholinergic enhancer AiE0743m_3xC2 driving nuclear SYFP2-H2B as a pool in both SCID and C57Bl6J, as this combination of enhancers should not have any significantly overlapping expression and was previously shown to produce transcriptional crosstalk in the cortex^6^ (**Fig. 4d**). In agreement with this, we observed rampant co-labeling of SYFP2 and mTFP1 that was worse in the cortex compared to striatum of C57Bl6J mice, with double positive cells accounting for 81% of all fluorescent cells in the cortex and 24% of fluorescent cells in the striatum (**Fig. 4e-f**). In SCID mice however, the number of co-labeled cells was reduced to 16% and 6% of fluorescent cells in the cortex and striatum, respectively (Fisher’s Exact Test, CTX ****P<0.0001, STR *P=0.0116), although the number of AiE0140h-mTFP1+ cells was also decreased in SCID mice. Taken together, these results further demonstrate DNA concatenation-dependent enhancer cross-activation can alter expression pattern characterization of enhancer AAVs.

## Discussion

With their compact size, cell type specificity, and conservation across species, enhancer elements are becoming increasingly more prevalent as tools for driving specific transgene expression *in vivo*. However, evaluating enhancer expression remains costly and time-consuming due to the lack of any efficient validation platforms. Using a combination of barcoded AAV vectors and single cell RNA sequencing, we aimed to increase screening throughput by adapting a multiplex approach. Instead, we identified multiple caveats including low AAV transcript capture and high technical and biological noise that prevented a complete characterization of all enhancer AAVs. These results demonstrate important caveats to consider when designing an enhancer AAV multiplexed screen.

Technical noise, derived from spurious ambient RNA or PCR chimeras, has been extensively documented in droplet-based RNA sequencing^23,29–31^ and is most likely the cause of low UMI counts. In contrast, biological noise is much more difficult to discern from true enhancer expression as these transcripts are produced by real enhancer-driven transcription (and why noise minimizing computer algorithms such as Cellbender failed in this context). These technical and biological sources of noise obscure multiplexed enhancer expression because they appear as real expression (**Fig. 4g**).

How can we minimize the effect of noise? Striking the right balance between enhancer AAV multiplex strength, composition, and size is key. A larger multiplex pool would distribute noise over a wider net, decreasing the amount of noise per enhancer. This effect was observed for the four enhancers included in both the six-plex (Figure 3) and twenty-plex (Figure 4), as the specificities were all higher in the twenty-plex (DLX2.0: 72% vs 97%, AiE0779m: 26% vs 36%, AiE0390m 75% vs 99%, AiE0410m: 75% vs 99%). Furthermore, the strength and completeness of labeling of each enhancer is also an important factor to consider. Signal from weak or sparse enhancers can be completely obscured by spurious signal arising from strong enhancers. We found this to be true of interneuron enhancers which exhibited some of the lowest expression specificities across multiplex pools.

Minimizing cross-expression by performing a multiplex screen in SCID mice was only partially successful. Although SCID mice reduced transcriptional crosstalk, it also had a profound and unwanted side effect of diminishing already low enhancer-driven expression levels (**Fig. S3**). This was especially evident in some cell types such as oligodendrocytes and parvalbumin interneurons, where reduced enhancer transcription significantly impacted calculated enhancer specificities. In other cell types, such as medium spiny neurons, reduced enhancer transcription did not affect the sensitivity of detection. This cell type discrepancy could be due to differences in expression of the SCID mouse LOF gene *Prkdc* or the exact mechanism of DNA repair (homologous vs non-homologous) across cell types^32^.

Another unexpected source of noise was from pooled AAV packaging, which we demonstrated was driven by rampant barcode swapping during AAV packaging. The observed rate of swapping (∼37%) aligns with the expected rate for homologous regions of this size, as reported in Lalanne et al. (co-submitted). Due to the large number of AAV screens using pooled packaging techniques and the widespread chimerism we observed, we are surprised that this phenomenon has only been documented now, as it has very likely contributed to high background noise in many such screens. Because of this, we caution against pooled AAV packaging which further limits the use of multiplexed enhancer screening. If pooled packaging is required, Lalanne et al. demonstrates possible swap-mitigation strategies.

With these caveats in mind, we believe enhancer AAV multiplexing can be useful for some screening applications. Two potential use cases would be a relatively small twenty-plex pool (packaged individually) used for rank ordering enhancers labeling similar cell types or screening enhancer optimization designs, both of which would be implemented to identify strong enhancers. Alternatively, a recent promising enhancer multiplex in human cell lines aimed to decouple enhancer signal from background noise by incorporating a second, constitutively expressed circular barcode within the screening construct^33^. Integrating these so-called “tornado barcodes” into our AAV screening vector would permit identification of cells that have been both transduced and express enhancer-derived transcripts. Regardless of exact strategy, major sources of biological and technical noise need to be identified and mitigated before multiplexed enhancer AAV screening can be correctly applied to classify enhancers with unknown specificity.

## Data and materials availability

10X Genomics sequencing data reported in this paper will be accessible in GEO (accession number to be determined). Allen institute whole mouse brain taxonomy (CCN20230722) is available at https://portal.brain-map.org/. All other data will be made available upon request.

## Acknowledgements

We wish to thank the Allen Institute founder, Paul G. Allen, for his vision and the NIH BRAIN Initiative Armamentarium for Precision Brain Cell Access (https://braininitiative.nih.gov/armamentarium). This research was supported by U.S. National Institutes of Health (NIH) BRAIN Initiative Armamentarium Grant UF1MH128339 (JTT, BPL, BT) and Allen Institute funding. We thank additional members of the core facilities and joint Brain Science shared resource teams including Laboratory Animal Services, Transgenic Colony Management Team, Animal Care Team, Viral Technology Team, Project Management Team, and Facilities Team. We also thank Nicholas Kamps-Hughes (Plasmidsaurus) for helpful discussions on long-read AAV sequencing.

## Author Contributions

<u>Conceptualization</u>: ACH, JKM, BPL, MG, J-BL, JTT. <u>Formal analysis</u>: ACH, JKM. <u>Investigation</u>: ACH, JKM, NT, AT, THP, DB, ABC, RC, RF, JG, JBG, NW. <u>Resources</u>: AT, THP, DB, ABC, RC, NPD, RF, KJ, JG, JBG, SK, RK, RAM, DN, NP, NVS, TZ, SY, KAS. <u>Supervision</u>: JKM, SY, KAS, ESL, BT, JTT, BPL, JS. <u>Writing</u>: ACH, JKM, BPL, MG.

## Declaration of interests

Authors JTT, BPL, BT, ESL, JKM are co-inventors on patent application PCT/US2021/45995 Artificial expression constructs for selectively modulating gene expression in striatal neurons. Authors JTT, BPL, BT are co-inventors on provisional patent application US 63/582,759 Artificial expression constructs for modulating gene expression in the basal ganglia. JKM and BPL declare equity in EpiCure Therapeutics, Inc. JS is a scientific advisory board member, consultant and/or co-founder of Cajal Neuroscience, Guardant Health, Maze Therapeutics, Camp4 Therapeutics, Phase Genomics, Adaptive Biotechnologies, Scale Biosciences, Sixth Street Capital, Prime Medicine, Somite Therapeutics and Pacific Biosciences.

Additional author affiliations

NW – University of Colorado School of Medicine, CO

## Supplemental Figures

**Figure S1:**
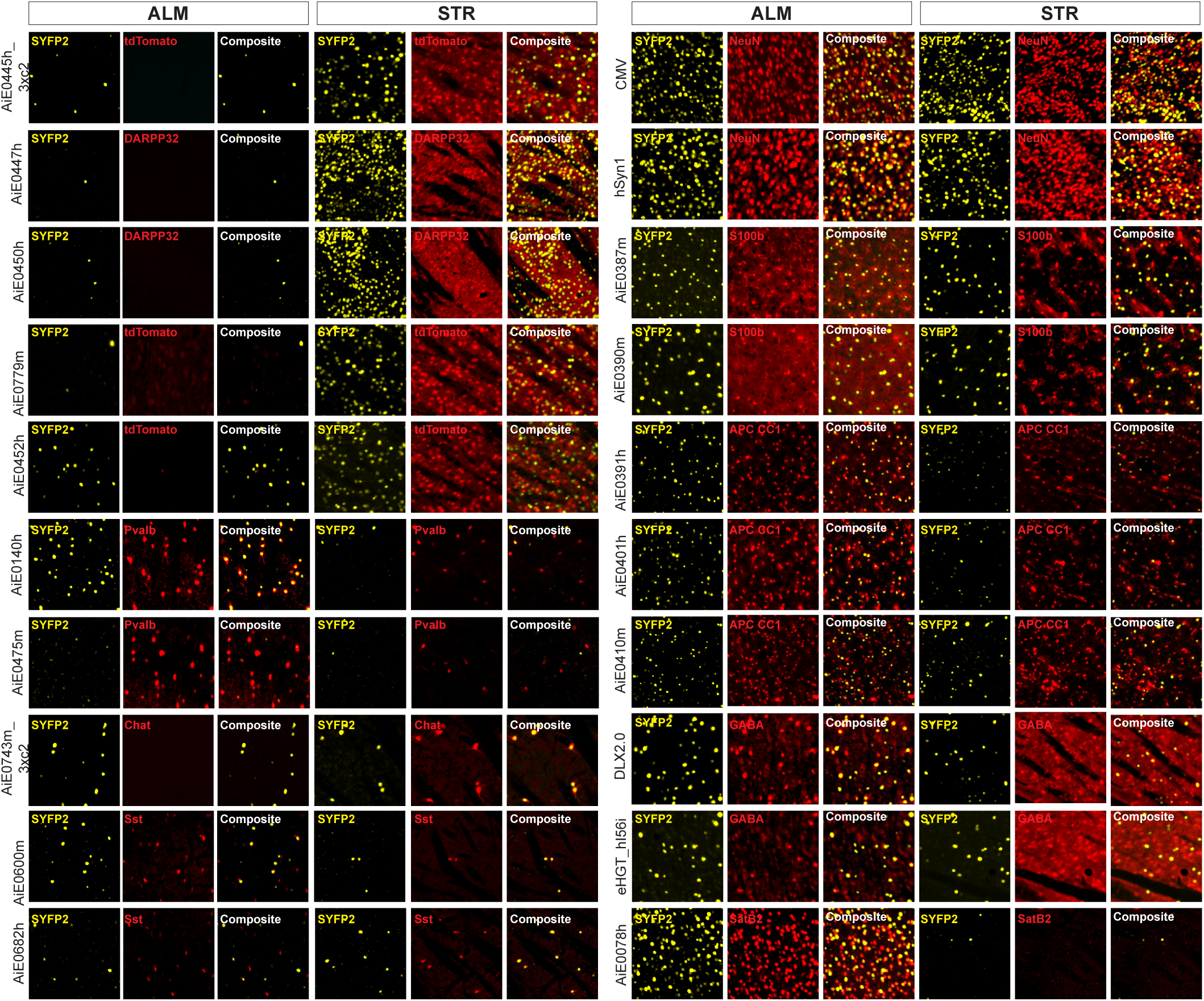
Enhancer specificities determined by IHC. IHC images of enhancer AAV SYFP2 expression following individual RO injections into adult mice. Enhancer specificity was evaluated by SYFP2 overlap with marker gene expression chosen based on published enhancer specificities. IHC images are from ALM (first and third column) and striatum (second and fourth column) and were used to generate the specificity data in Figure 1g.

**Figure S2:**
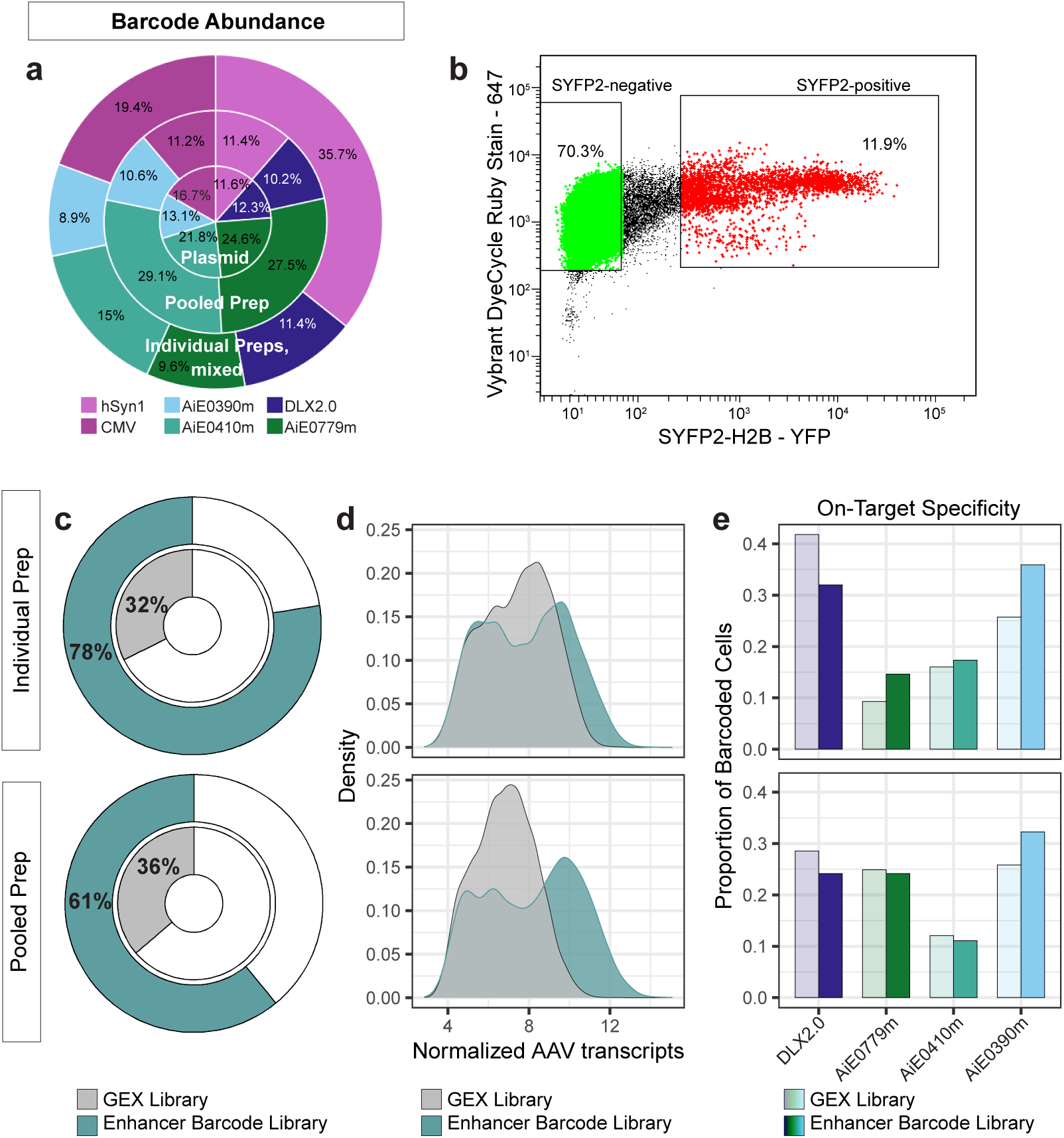
Generation of enhancer barcode library is necessary for detection of enhancer AAV transcripts. **a)** Barcode abundance (shown as % of reads) for individual preps mixed at equal proportions (outer ring), pooled prep (middle ring) and combined plasmids (inner circle) determined by Amplicon-EZ short read sequencing prior to mouse administration. **b)** Example of gating strategy from whole cells isolated from striatum of mouse injected with individual preps, mixed. **c-d)** Comparison of SYFP2-positive cells with barcoded transcripts in individual (top) vs pooled (bottom) AAV preps. **c)** Percent of cells in either the GEX library or AAV barcode library containing transcripts with enhancer barcodes. **d)** Distribution of transcripts per cell (UMIs) across all six AAVs. **e)** Proportion of enhancer barcoded cells that are on-target pre-filtering.

**Figure S3:**
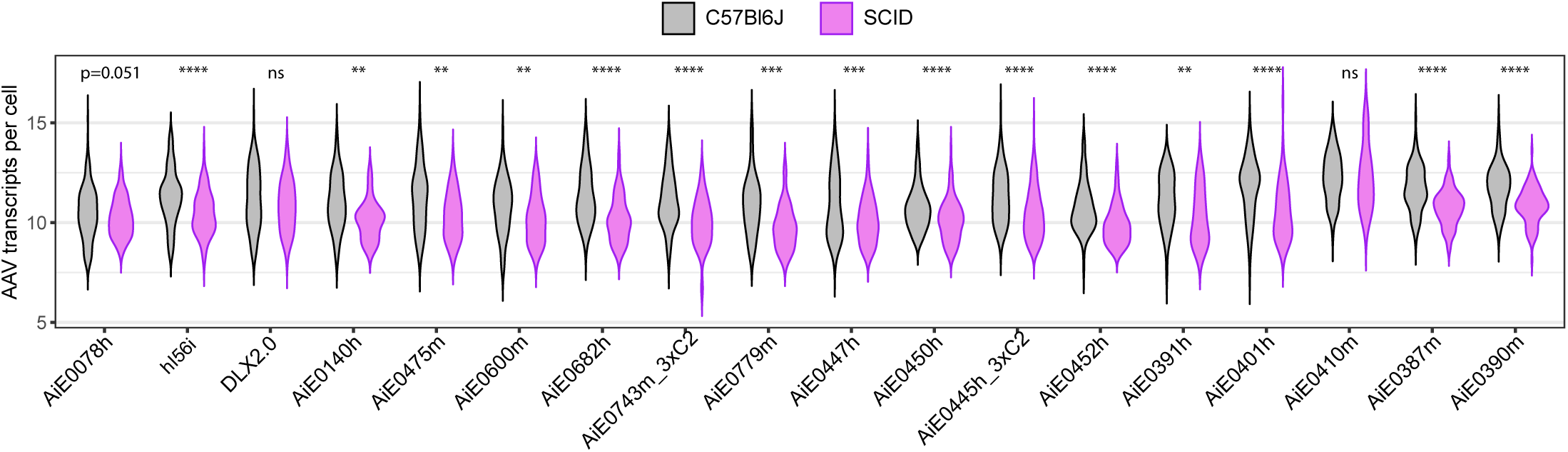
Reduced enhancer-driven transcription in SCID mice. Violin plots of the normalized AAV transcript (UMI) count distribution for each enhancer, grouped by mouse line. Normality was evaluated using the Shapiro-Wilk test. Statistical significance was assessed using a Mann-Whitney U test for each enhancer. Stars indicate significance levels: ****p<0.0001, ***p<0.001, **p<0.01, *p<0.05.

## Methods

### Experimental model and subject details

All experiments were performed in accordance with the Allen Institute Institutional Animal Care and Use Committee (IACUC) (protocols 2002, 2105, 2301, and 2406). All mice are on a C57BL6J background and were group-housed (max 5 mice/cage) on a twelve-hour light/dark cycle with ad libitum access to food and water. C57Bl6J (Stock 000664) and *Prkdc*^scid/scid^ (SCID mice, Stock 001913) mice were purchased from Jackson Laboratories. Drd1a-tdTomato line 6 hemizygous mice ^34^ were bred in house. Mice were injected between ages P28-P56.

### Design and cloning barcoded enhancer AAVs

The 8bp barcode strategy was derived from Guo et al., 2019^35^ and all twenty barcodes used in this study were designed to have a Hamming Distance >2. Barcodes were ordered as single-stranded forward and reverse compliment oligonucleotides from Integrated DNA Technologies (IDT) with 20bp overlapping homology regions on either side of the XhoI site in the vector rAAV-hSyn1-SYFP2-10aa-H2B-WPRE3-bGHpA (CN1839, Addgene 163509). Each lyophilized oligonucleotide stock was resuspended in water to 100µM and then combined and diluted 1:500 with its pair. Each pair was annealed at 100°C for 5 minutes and slowly cooling to room temperature over 30 minutes. The annealed oligonucleotides were cloned into the vector using In-Fusion HD Cloning Kit (Takara 639650). Enhancers plus minimal promoters were subcloned from plasmids in house into the barcoded vectors using MluI/AgeI restriction enzyme sites and T4 Quick Ligase (NEB M2200L). Every enhancer was paired with a single, unique 8bp barcode sequence.

| Enhancer/Promoter | Barcode 5' → 3' |
| --- | --- |
| hSyn1 | TGCTATAT |
| AiE0390m | GTTGGCTA |
| AiE0410m | AGTTTAGG |
| CMV | GGTTCACA |
| DLX2.0 | TAGAAAGC |
| AiE0779m | TCGCCTTA |
| AiE0387m | CTAGTACG |
| AiE0401h | TTCTGCCT |
| AiE0391h | GCTCAGGA |
| hI56i | CATGCCTC |
| AiE0078h | GTAGAGAT |
| AiE0140h | AACTGTAG |
| AiE0475m | GACGTGAC |
| AiE0452h | TATACGAG |
| AiE0445h_3xC2 | ATGCATGC |
| AiE0450h | TCACCGAT |
| AiE0447h | CGGCTAAT |
| AiE0682h | GATCCACC |
| AiE0600m | AGCAATTA |
| AiE0743m_3xC2 | CCGGAATT |

Infusion reactions were transformed into chemically competent *Stbl3* E. coli (ThermoFisher C737303) and individual colonies were selected from 100 µg/mL carbenicillin plates. Plasmids were maxiprepped (Qiagen 12162) and verified using Azenta Sanger sequencing.

### AAV Production

All plasmids used in this study were packaged into the AAV capsid PHP.eB^17^ using a small-scale crude prep packaging method as described previously^6,10^. The cells were seeded at 2 × 107 cells per 15-cm dish to achieve ∼70% to 80% confluency before transfection. Cells were maintained in DMEM (Thermo Fisher Scientific Cat#1995-065) with 10% Fetal Bovine Serum (FBS; VWR Cat#89510-184) and 1% Antibiotic-Antimycotic solution (Sigma Cat#A5955). Small-scale crude AAV preps were generated by triple transfecting 15 µg ITR plasmid, 15 µg AAV PHP.eB capsid plasmid, and 30 µg pHelper (Cell Biolabs) in 1.35 ml of OptiMem (Thermo Fisher Scientific Cat#51985-034), which was then supplemented with 150 µl of 1 mg/mL Polyethylenimine (PEI; Polysciences Cat#23966), incubated for 10 minutes and then added to a single 15 cm plate of fully confluent cells.. For pooled packaging, plasmids were combined at equal ratios for a total of 15 µg. At one day post-transfection medium was replaced with low serum (1% FBS), and after 3 days the cells and supernatant were collected, freeze-thawed 3x to release AAV particles, treated with benzonase nuclease (Millipore Sigma E8263-25KU) for 30 minutes to degrade free DNA, then clarified (3000g × 10min) and concentrated to approximately 150 µL by using an Amicon Ultra-15 centrifugal filter unit (Sigma Cat # UFC910024) at 5000xg for 30-60 min. Viral prep titers were assessed using digital droplet PCR (BioRad QX200 system) using primers against AAV2 ITRs and typically resulted in around 100µL of 1-5E13 vg/mL per virus.

To determine barcode representation in combined or pooled AAV preps, viruses were diluted 1:1000 in water, denatured at 100°C for 5 minutes, and PCR amplified using KAPA HiFi Mastermix (Roche 7958935001) with forward primer 5’ AACTCATCGCCGCCTGCCTTG 3’ and reverse primer 5’ ACAGTGGGAGTGGCACCTTC 3’ for 30 cycles. Following amplification, reactions were purified using 0.6x and 0.9x SPRI beads, resuspended in 25µL Buffer EB, and sequenced using Azenta Amplicon-EZ sequencing service. Sequencing reads were visualized using R package ShortRead and barcodes were identified using string_detect from R package stringr. Percent barcode representation was calculated as number of reads with specific barcode*X* divided by total reads with any valid barcode.

To check for recombination during pooled packaging, AAV preps were sequenced using Oxford Nanopore long read sequencing using the AAV Genome Sequencing service from Plasmidsaurus. Raw sequencing reads were mapped against the six AAV genomes using minimap2 and split by which plasmid they mapped to. Partial reads were discarded, and barcodes were identified using string_detect from R package stringr. Percent valid enhancer-barcode pairs were calculated by valid enhancer-barcode pair divided by total enhancer-barcode pairs and multiplied by 100 to obtain a percentage.

### Retro-orbital mouse injections

PHP.eB-packaged AAVs were first diluted either individually (5E11 vg) or in a pool (6E11-1E12 total vg) for the multiplex experiments in 100uL of sterile 1xPBS. Adult P40-P56 male mice were injected with the diluted AAVs into the retro-orbital sinus after isoflurane anesthesia. Tissue was harvested 4-5 weeks post-injection for all experiments.

### Mouse regional brain dissections

Mice were anesthetized with Avertin and perfused transcardially with either ice-cold Cutting Buffer containing 110mM NaCl, 2.5mM KCl, 10mM HEPES, 7.5mM MgCl2, 25mM glucose, and 75mM sucrose for cell isolations or 1xPBS for nuclear isolations. Following perfusions, brains were extracted and placed in a petri dish containing Cutting Buffer (cells) or 1xPBS (nuclei) to generate 2mm coronal slices using a stainless-steel brain matrix (Stoelting 51386). Slices containing ALM or STR were transferred to a second petri dish with Dissociation Buffer containing 82mM Na2SO4, 30mM K2SO4, 10mM HEPES, 10mM Glucose, and 5mM MgCl2 (cells) or 1xPBS (nuclei). ALM and STR were isolated from the coronal sections using a disposable safety scalpel (VWR 21909-670). Tissue pieces were then processed immediately for cell isolations or placed in 1.5mL Eppendorf tubes, flash frozen in a dry ice-ethanol bath and stored at −80°C for nuclei isolations.

### Single cell isolations

Cell suspensions were prepared as previously described^36^. After dissection, ALM, or STR were placed separately in 10mL conical tubes with 5mL of room temperature Enzyme Buffer containing 3mg/mL Protease XXIII (Sigma-Aldrich P5380) and 10 units/mL of Papain (Worthington LK003150) dissolved in Dissociation Buffer. Submerged slides were immediately transferred to a 34°C water bath for 1 hour (STR) or 2 hours (ALM). Following digestion, tissue was placed directly on ice and the supernatant was carefully removed and replaced with 10mL Stop Solution containing 1mg/mL Trypsin Inhibitor (Sigma-Aldrich T6522), 2mg/mL BSA (Sigma-Aldrich A2153), and 1mg/mL Ovomucoid Protease Inhibitor (Worthington LK003150) dissolved in Dissociation Buffer. Tissue chunks were triturated using four fire-polished glass Pasteur pipets with successively smaller bore holes while avoiding bubbles until a homogenous solution was obtained. Triturated tissue was centrifuged at 300xg for 10min at 4°C, supernatant decanted, pellet resuspended in 5mL of Stop Solution, and then centrifuged again at 300xg for 10min at 4°C. Following centrifugation, the supernatant was decanted once more and the cell pellet was resuspended in 500uL-1mL of Dissociation Buffer with 0.1% BSA, 1:500 DAPI (ThermoFisher 62248), and 1:500 Vibrant DyeCycle Ruby Stain (ThermoFisher V10309). The resulting single cell suspensions were filtered using a pre-wet 70µm filter (Miltenyi Biotec 130-098-462) and incubated in the dark at 4°C for 30 minutes prior to flow cytometry.

### Single nucleus isolations

Single nucleus isolations were prepped as described previously ^19^. In brief, 250mL of Nuclei Isolation Media (pH=8) containing 250mM sucrose, 10mM Tris Buffer (pH=8), 25mM KCl, 5mM MgCl_2_ was prepared ahead of time and placed at 4°C for up to 2 weeks. Next, 2mL glass dounces with their accompanying A and B pestles (Sigma-Aldrich D8938) were placed in RNase Away solution (VWR 53225-514) overnight at room temperature. The following day, the dounces were rinsed thoroughly with MilliQ water and air dried. Dounces were placed on ice and filled with 1.5mL of Homogenization Buffer containing 0.1mM DTT (Promega P1171), 1x Protease Inhibitor Cocktail (Promega G6521), 0.2 U/µL RNasin Plus RNase Inhibitor (Promega N2615), 0.1% Triton X-100 dissolved in 0.22um filtered Nuclei Isolation Media. Previously frozen ALM or STR tissue punches were retrieved from −80°C and transferred to the Homogenization Buffer-filled dounces. Tissue was homogenized 15-20x with Pestle A, followed by 10-15x with Pestle B and then filtered through a 70µm filter (Miltenyi Biotec 130-098-462) on top of a 15mL conical tube. An additional 3mL of Homogenization Buffer was added and then the tissue was centrifuged at 900xg for 10min at 4°C. Following centrifugation, the supernatant was discarded, and the nuclei pellet was resuspended in 1mL of Blocking Buffer containing 1x PBS (pH=7.4), 1% BSA (Sigma-Aldrich A9576), 0.2 U/µL RNasin Plus RNase Inhibitor, and transferred to a 1.5mL Eppendorf tube. DAPI (0.1 µg/mL) and 1:500 mouse monoclonal anti NeuN conjugated to PE (Sigma-Aldrich FCMAB317PE) was added to the tube and incubated at 4°C for 30min in the dark, then centrifuged at 400xg for 5 minutes at 4°C. Supernatant was decanted, and nuclei pellet was resuspended in 500uL of Blocking Buffer and filtered through the cap of a 5mL FACS tube (Fisher Scientific 08-771-23) on ice prior to flow cytometry.

### Flow cytometry

Cell and nuclei suspensions were sorted for SYFP2 expression using a 130µm nozzle on a BD FACSAria III at a flow rate of <2000 cells/nuclei per second on purity mode. Live cells were selected for by a DAPI-negative, Ruby-positive gate and intact nuclei by a DAPI-positive gate. A goal of 50-100,000 SYFP2-positive and SYFP2-negative cells and nuclei were collected in chilled 5mL FACS tubes containing 300uL Dissociation Buffer with 0.1% BSA (cells) or Blocking Buffer (nuclei). Following FACS, BSA was added to the sorted nuclei for a final concentration of 2%, centrifuged at 500xg for 5 minutes at 4°C, and all but ∼50uL of supernatant above the pellet was decanted. The nuclei were resuspended in Freezing Media containing 1xPBS, 1% BSA, 10% DMSO (Millipore-Sigma 276855), and 0.2 U/µL of RNasin Plus RNase Inhibitor at a 1:1 ratio and frozen at −80°C until loading on the 10X Genomics v3.1 chip (1000268, 10x Genomics) later. The sorted cells were centrifuged at 300xg for 10min at 4°C, supernatant decanted, and the pellet resuspended in 80-100uL of Dissociation Buffer with 0.1% BSA on ice. Cell concentrations were estimated using a disposable hemocytometer (Thomas Scientific 1190G82) on a Nikon Ti-Eclipse microscope and were immediately loaded on the 10X v3.1 chip for insertion into the 10X Genomics Chromium controller.

### 10X Genomics, custom AAV barcode library generation, and sequencing

Previously sorted nuclei were removed from the −80°C and allowed to thaw on ice. 100uL of Nuclei Wash and Resuspension Buffer containing 1% BSA, 0.2U/µL Rnasin Plus RNase Inhibitor in 1xPBS was added to the nuclei and then centrifuged at 400xg for 5 minutes at 4°C. Following centrifugation, the supernatant was removed, and the nuclei pellet was resuspended in <50µL of Nuclei Wash and Resuspension Buffer. Nuclei concentrations were estimated using a disposable hemocytometer and concentration was adjusted to 1000 nuclei/uL using Nuclei Wash and Resuspension Buffer. Approximately ∼16,000 cells or nuclei were loaded on the 10X Genomics v3.1 chip (1000268, 10x Genomics) for all experiments. For GEX library generation, the manufacturer’s instructions for cell capture, barcoding, reverse transcription, cDNA amplification and library construction were followed. Libraries were sequenced on the Illumina NovaSeq 6000 or NovaSeq X with a target read depth of 125,000 reads per cell/nuclei.

To generate custom libraries for the AAV barcoded transcripts, 10ng of the amplified cDNA from each sample was first amplified in a 100uL reaction containing 50µL KAPA Hifi Mastermix (Roche 7958935001), 0.5µL 100µM forward WPRE3 primer 5’ AACTCATCGCCGCCTGCCTTG 3’ and 0.5µL 100uM reverse Read 1 primer 5’ CTACACGACGCTCTTCCGATCT 3’ and amplified using a thermocycler with the following conditions: 98°C for 5 minutes initial denaturation, then 8x cycles of 95°C for 15 seconds, 60°C for 30 seconds, 72°C for 20 seconds, and lastly 72°C for 30 seconds. The reaction was purified using 0.7X and 0.9X SPRI beads (Beckman Coulter B23318) and then resuspended in 40µL Buffer EB (Qiagen 19086).

Sequencing adapters and indexed were added using 20µL of product from the first PCR along with 0.5µL of 100µM nested forward primer 5’ AATGATACGGCGACCACCGAGATCTACACNNNNNNNNNNACACTCTTTCCCTACACGACGCTCTTCCGATCT 3’, 0.5µL of 100uM reverse primer 5’ CAAGCAGAAGACGGCATACGAGATNNNNNNNNNNGTGACTGGAGTTCAGACGTGTGCTCTTCCGATCTACTGACAATTCCGTGGCTCG 3’, and 50µL KAPA HiFi MasterMix. Reaction was amplified using 12 cycles, purified using 0.7x and 0.9x SPRI beads, resuspended in 25µL Buffer EB, and checked for purity and concentration using the High Sensitivity Kit (Agilent 067-4626) on a Bioanalyzer. Custom libraries were pooled prior to sequencing on the Illumina NextSeq 2000 with a target read depth of ∼5,000 reads per cell.

### Sequencing data processing and QC filtering

A custom reference was created using the 10x Genomics Cellranger mkref to add the AAV vector sequence SYFP2-H2B-WPRE3-[barcode]-bGHpA to the mouse reference transcriptome (M21, GRCm38.p6). Sequencing fastq read files were aligned to the custom reference using the 10x Genomics Cell Ranger pipeline (version 6.1.1) with default parameters. AAV barcode UMI counts were identified in the AAV barcode library fastqs using the shell scripting tool BarCounter ^37^ adapted for the vector genome. The pattern used to identify the AAV barcodes was 5’ AG(BC) 3’.

High-quality cells and nuclei were selected for using stringent QC filtering ^32,38^ using the following filters: mitochondrial gene percentage (<5% for nuclei, <20% for cells), total genes detected (>1000 for nuclei, >2000 for cells) and a doublet score of >0.3 using DoubletFinder ^39^. For multiplexed libraries, cells that did not contain AAV transcripts were also discarded.

Following QC filtering, cell and nuclei gene x cell count matrices were normalized using logCPM and mapped using map_by_cor (both part of R package scrattch.hicat) onto the CCN20230722 whole mouse brain taxonomy ^32^. Cells or nuclei were discarded if they did not map to cell types in the corresponding region (either ALM or STR) and if they did not map with at least 80% confidence to the reference taxonomy.

AAV expression was normalized by using logCPM on the AAV barcode x cell matrix produced from BarCounter appended to the GEX matrix. Normalized UMI counts were visualized using a density plot. For cell types exhibiting a bimodal distribution in enhancer AAV barcodes, UMI counts below the antimode were discarded. Z-scores were calculated for each enhancer AAV across cell types using the equation (C_E_ – C_P_)/ sd(C_E_), where C_E_ = number of enhancer-labelled cells in cell type and C_P_ = number of predicted cells per cell type. Number of predicted cells were calculated by (T_E_ x T_C_)/ T_L_, where T_E_ = total number of enhancer-labelled cells, T_C_ = total number of cells in cell type and T_L_ = total number of cells in library. Cell types with negative Z-scores (de-enriched populations) were discarded. The remaining cell populations were used to determine enhancer on-target cell type specificities which was calculated as number of cells in on-target cell types divided by total number of remaining enhancer-labelled cells multiplied by 100 to obtain a percentage.

### Immunohistochemistry

Mice injected with enhancer AAVs were anesthetized with Avertin and perfused transcardially with ice-cold phosphate-buffered saline (PBS) followed by 4% paraformaldehyde (PFA). All enhancer AAVs were injected into C57Bl6J mice except for D1 MSN enhancer AiE0779m and D2 MSN enhancers AiE0445h_3xC2 and AiE0452h which were injected in Drd1a-tdTomato line 6 hemizygous mice. The brain was extracted and placed in 4% PFA overnight, then transferred to 30% sucrose in PBS for 2-3 days. Brains were cut in half and mounted on a thin block of OCT cryo-compound (Tissue-Tek # 4583) on a frozen sliding microtome for generation of 30µm sagittal sections. Sections were stored in PBS with 0.02% sodium azide at 4°C. For IHC, stored slices were washed 3x in PBS and then placed in blocking buffer containing 5% normal goat serum, 0.1% TritonX-100 in PBS, and primary antibody(ies) at 4°C overnight. All primary antibodies were diluted to a 1:1000 concentration except for mouse SATB1 + SATB2 which was used at 1:250. The following day, sections were washed 3x in wash buffer containing 0.1% TritonX-100 in PBS and then placed in blocking buffer with appropriate secondary antibodies diluted 1:1000 for 2 hours at room temperature. Secondary antibodies included Alexa Fluor 488 Goat anti Chicken (ThermoFisher A11039), Alexa Fluor 555 Goat anti Rabbit (ThermoFisher A32732), Alexa Fluor 555 Goat anti Mouse IgG1 (ThermoFisher A21127), and Alexa Fluor 555 Goat anti Mouse IgG2b (ThermoFisher A21147). Sections were washed 3x in wash buffer and then mounted onto microscope slides using Prolong Gold and imaged at 10x on a Nikon Ti-Eclipse or Nikon Ti-Eclipse 2 epifluorescent microscope.

Specificity of enhancer activity for all enhancers (except D2 MSN enhancers) for the target cell subclass or type was quantified as reporter and marker antibody double positive neuron count divided by total reporter antibody positive neuron count and multiplied by 100 to obtain a percentage.

Specificity for D2 MSN enhancers injected into Drd1a-tdTomato mice was determined by quantifying non-overlapping populations (1 - reporter and marker antibody double positive neuron count divided by total reporter Ab positive neuron count and multiplied by 100 to obtain a percentage). An important caveat to this strategy is it would include any labeling of striatal interneuron populations as on-target. However, both our mapping (figure 4) and published validations (Hunker, Wirthlin et al., 2024) suggest interneuron labeling is minimal for these enhancers.

| Antibodies | Source | Identifier |
| --- | --- | --- |
| Chicken polyclonal anti GFP | Aves | GFP-1010 |
| Rabbit polyclonal anti NeuN | Millipore | ABN78 |
| Mouse monoclonal anti S-100 (β subunit) | Sigma | S2532 |
| Mouse monoclonal anti APC [CC-1] | Abcam | ab16794 |
| Rabbit polyclonal anti GABA | Sigma | A2052 |
| Mouse anti SATB1 + SATB2 antibody | Abcam | ab51502 |
| Rabbit polyclonal anti Parvalbumin | Swant | PV27a |
| Mouse monoclonal anti DARPP-32 | BD Biosciences | 611520 |
| Rabbit polyclonal anti RFP | Rockland | 600-401-379 |
| Rabbit polyclonal anti somatostatin-28 | Peninsula Labs | T-4547 |
| Mouse monoclonal anti ChAT | Atlas | CL3169 |
| Rat anti mTFP | CancerTools | 155264 |

